# Prognostic association of immunoproteasome expression in solid tumours is governed by the immediate immune environment

**DOI:** 10.1101/2022.08.30.505767

**Authors:** Rahul Kumar, Bhavya Dhaka, Sarthak Sahoo, Mohit Kumar Jolly, Radhakrishnan Sabarinathan

## Abstract

Induction of immunoproteasome (IP) expression in tumour cells can enhance antigen presentation and immunogenicity. Recently, overexpression of IP genes has been associated with better prognosis and response to immune checkpoint blockade (ICB) therapies in melanoma. However, the extent of this association in other solid tumour types and how that is influenced by tumour cell-intrinsic and cell-extrinsic factors remains unclear. Here, we address this by exploring the gene expression patterns from available bulk and single-cell transcriptomic data of primary tumours. We find that IP expression positively correlates with the constitutive proteasome (CP) across multiple tumour types. Furthermore, tumours with high IP expression exhibit cytotoxic immune cell infiltration and upregulation of interferon-gamma and TNF-α pathways in tumour cells. However, the association of IP expression with overall survival (in TCGA cohort) and response to ICB therapy (in non-TCGA cohorts) is tumour-type specific and is greatly influenced by immune cell infiltration patterns. This emphasises the need for considering immune cell infiltration patterns, along with IP expression, to be used as a prognostic biomarker to predict overall survival or response to ICB treatment in solid tumours, besides melanoma.

## 1. Introduction

Proteasomes are an essential component of the ubiquitin-proteasome system that maintains protein homeostasis by degrading unwanted or misfolded proteins into smaller peptides. These peptides are then processed by the antigen-processing machinery and presented on the cell surface by the major histocompatibility complex (MHC) class I molecules for immune surveillance (1,2). The proteasome complex (26S) constitutes one catalytic core (20S) and two terminal regulatory subunits (19S). The proteasome exists primarily in two forms: constitutive proteasome (CP) and immunoproteasome (IP) (and sometimes in an intermediate form as well) (3–5). The CP is ubiquitously expressed across all nucleated cell types, and its 20S core contains three catalytic cleaving subunits: β5/PSMB5 (chymotrypsin-like), β1/PSMB6 (caspase-like), and β2/PSMB7 (trypsin-like). Whereas in the IP (or 20Si), which are predominantly expressed in immune cells (e.g., antigen-presenting cells), the above three subunits are replaced by β5i/PSMB8 (chymotrypsin-like), β1i/PSMB9 (chymotrypsin-like), and β2i/PSMB10 (trypsin-like) (2,4,6).

In non-immune cells, expression of IP genes can be induced by exposure to specific proinflammatory cytokines such as interferon-gamma (IFN-γ) and tumour necrosis factor alpha (TNF-α) under stress or inflammatory conditions (6). Following that, replacement of CP with IP at the protein level is mediated by proteasome maturation protein (POMP), which favours the incorporation of the IP subunit due to its faster assembly (∼4-fold faster) than CP (7–9). The IP has been shown to generate a higher immunogenic spectrum of peptides due to its altered cleavage-site preferences (∼32% unique sites as compared to CP) and also a higher number of spliced peptides (81%) than CP (65%) (2,10,11). This can lead to enhanced antigen presentation and thereby trigger the immune response. Genetic alterations and expression dysregulation of single or multiple subunits of IP have been associated with reduction of MHC class I expression (in non-immune cells), loss of B-cells, defects in T-cell development and T-helper cell differentiation, cytokines dysregulation, and reduced numbers of antigen-specific CD8+ T cells (12–15).

Although the role of IP expression and its prognostic association has been extensively studied in hematological malignancies (16,17), minimal studies have explored it in the context of solid tumours (see review (18)). A recent study in melanoma has shown that overexpression of IP subunits (PSMB8 and PSMB9) was associated with improved survival and better response to anti-CTLA-4 and anti-PD-1 immunotherapies (19). Similarly, in breast cancer, interferon-gamma mediated higher expression of IP was associated with better prognosis (20). In non-small-cell lung cancer (NSCLC), patients with higher PSMB9 expression showed better response to anti-PD-1 and progression-free survival (21). However, the extent of this association in other solid tumour types, and the influence of tumour cell-intrinsic and cell-extrinsic factors, remains unclear. Thus, in this study, we set out to address the following questions: (a) how are CP and IP gene expression (dys)regulated in tumours (that is, when the IP genes are induced, what happens to the CP expression); (b) what proportion of the cells are expressing CP and IP within the tumour; (c) what tumour cell-intrinsic and cell-extrinsic factors influence the expression of CP and IP; and (d) how does that affect the prognostic association and response to immune checkpoint blockade (ICB) therapies.

Through gene expression analysis of bulk and single-cell transcriptomic data of primary tumour samples (from TCGA and other studies, respectively), we show that (a) the expression of IP and CP was positively correlated in multiple tumour types as compared to normal tissues; however, the expression of IP was relatively lower than CP in both tumours and normals; (b) the expression of IP was not only predominant in immune cells infiltrating the tumours, but also in the subset of tumour epithelial cells (around 33% of the total epithelial cells). Notably, the tumour epithelial cells located in the tumour border expressed higher IP and CP than those in the tumour core regions; (c) the expression of IP was positively correlated with the upregulation of hallmark pathways such as interferon-gamma response, interferon-alpha response, IL-6-JAK/STAT3 signalling, and reactive oxygen species (ROS) pathway; while it was negatively correlated with upregulation of Wnt/beta-catenin signalling, TGF-β signalling and hedgehog signalling; (d) the enrichment of cytotoxic immune infiltrating cells (such as activated CD8+ T cells, gamma-delta T cells and natural killer cells) in the tumour microenvironment was positively correlated with higher expression of IP in tumour cells, whereas for eosinophils and T-helper cells, it was negatively correlated; (e) the association of IP expression with overall survival and response to ICB therapies was tumour-type specific and was greatly influenced by immune-cell infiltration patterns (pro-or anti-tumorigenic). Taken together, these results suggest that the expression of IP in tumour cells can be influenced by both cell-intrinsic and cell-extrinsic factors. Thus, the expression of IP subunits, combined with tumour immune cell infiltration patterns, can be used as a better predictor of prognosis and response to ICB therapies in solid tumours besides skin melanoma.

## 2. Results

### 2.1 Immunoproteasome expression is higher in tumours than normal tissues

To study the expression patterns of the constitutive proteasome (CP) and immunoproteasome (IP), we first compared the average expression scores of genes encoding the catalytic subunits of CP (PSMB5, PSMB6, and PSMB7) and IP (PSMB8, PSMB9, and PSMB10) in 9,491 TCGA tumour samples, across 33 cancer types (see Methods, Table 1, Supplementary Table 1). For most solid tumour types, we observed a significantly (*P*<0.05) higher expression of CP as compared to IP (Figure 1A). A similar trend was observed when controlled for tumour purity (>70%) to reduce the confounding effects from the stromal and immune microenvironment (Supplementary Figure 1). However, in tumours of hematopoietic origin – diffuse large B-cell lymphoma (DLBC) and acute myeloid leukemia (LAML) – a higher IP-to-CP expression was observed, which can be explained by the innate IP expression in hematopoietic cells. As compared to CP, the expression of IP was highly variable (large interquartile ranges) within each tumour type (Figure 1). This could be due to the influence of tumour cell-intrinsic and/or cell-extrinsic factors (see Sections 2.3 and 2.4). Nevertheless, we observed a positive correlation between CP and IP expression in multiple cancer types, suggesting that factors influencing induction of IP expression also have a positive effect on CP expression in tumours (Figure 1A).

**Figure 1:**
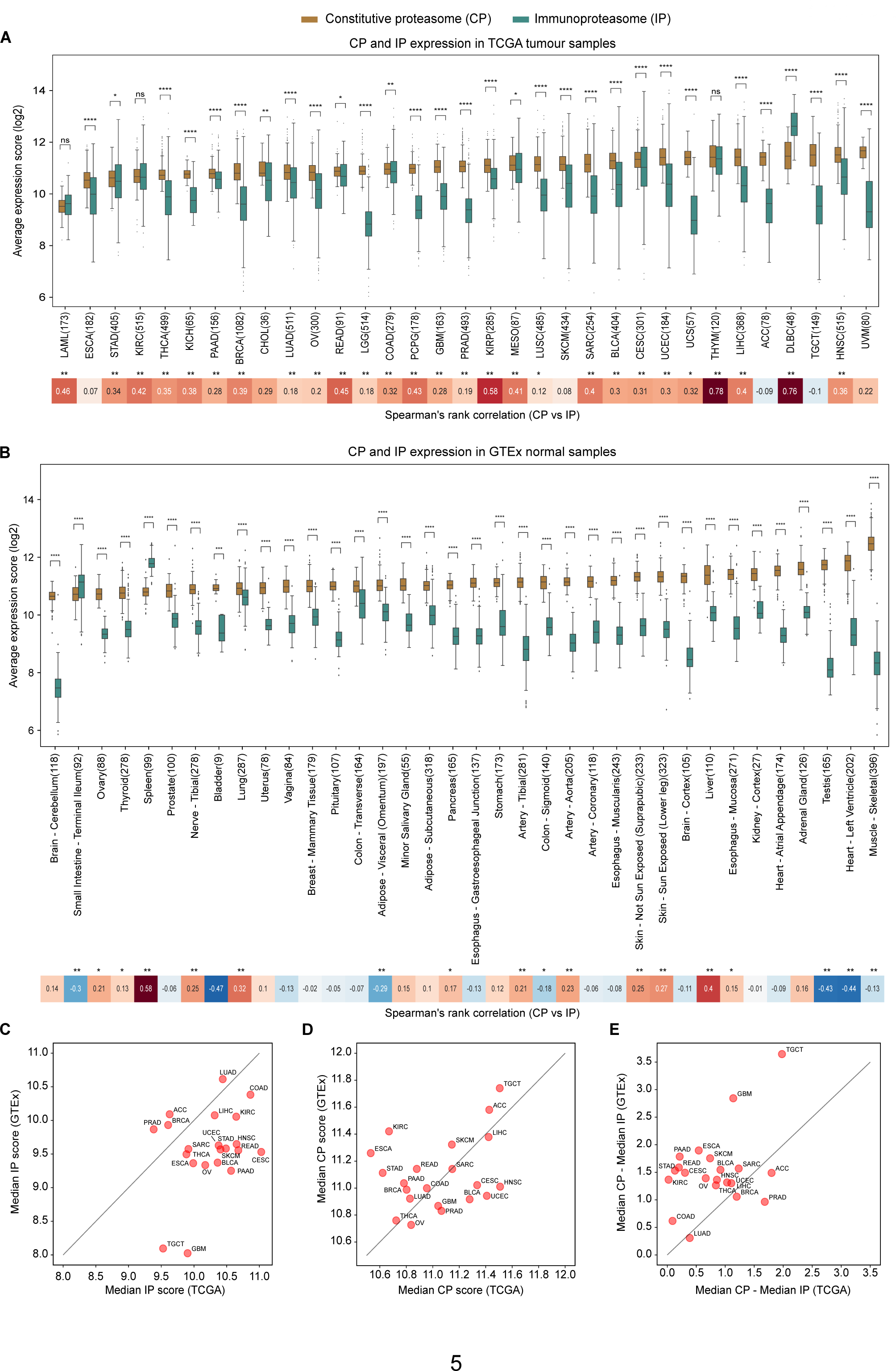
Comparison of CP and IP expression in tumours and normal tissues. **A.** The box plot shows the distribution of average expression of constitutive proteasome (CP) and immunoproteasome (IP) genes at the sample level across 33 different tumour tissues from TCGA. The x-axis represents the tumour types (with the number of samples) and the y-axis represents the average expression level of CP (PSMB5, PSMB6 and PSMB7) and IP (PSMB8, PSMB9 and PSMB10) genes (see Methods). See Table 1 for the tumour type full form for the abbreviations shown on x-axis. In each boxplot, the horizontal middle line indicates the median, the height of the shaded box indicates the interquartile range (IQR), and the whiskers indicate 1.5 x IQR. The P-value shown at the top, comparing IP and CP expression distributions at each tumour type, was computed using the Mann-Whitney U test (two-sided) and the significance level was represented as: **** *P* <= 0.0001, *** 0.0001 < *P* <= 0.001, ** 0.001 < *P* <= 0.01, *0.01 < *P* <= 0.05 or ns - non-significant (*P* > 0.05). The heatmap at the bottom represents Spearman’s rank correlation between average expression of CP and IP at the sample level for each tumour type. The tumour types that showed significant correlation were highlighted with the asterisks symbol on the top (** *P* <= 0.01 and * 0.01 < *P* <= 0.05). **B.** Same as A, but for the normal tissues from GTEx. The x-axis represents the normal tissues (with the number of samples) and the y-axis represents the average expression level of CP (PSMB5, PSMB6 and PSMB7) and IP (PSMB8, PSMB9 and PSMB10). **C.** Comparison of the IP expression level in TCGA tumours with respect to its matched normal tissues from GTEx. The x-axis and y-axis represent the median of the average expression of IP in tumours (as shown in Figure 1A) and in the matched normal tissues in GTEx (as shown in Figure 1B), respectively. **D.** Same as C, but for the CP. **E.** Comparison of differences in CP and IP expression level in TCGA tumours with respect to its matched normal tissues from GTEx. The x-axis shows the difference in the median of average expression of CP and IP expression value in TCGA tumours and the y-axis represents the same difference in the corresponding normal tissue from GTEx.

**Table 1:**
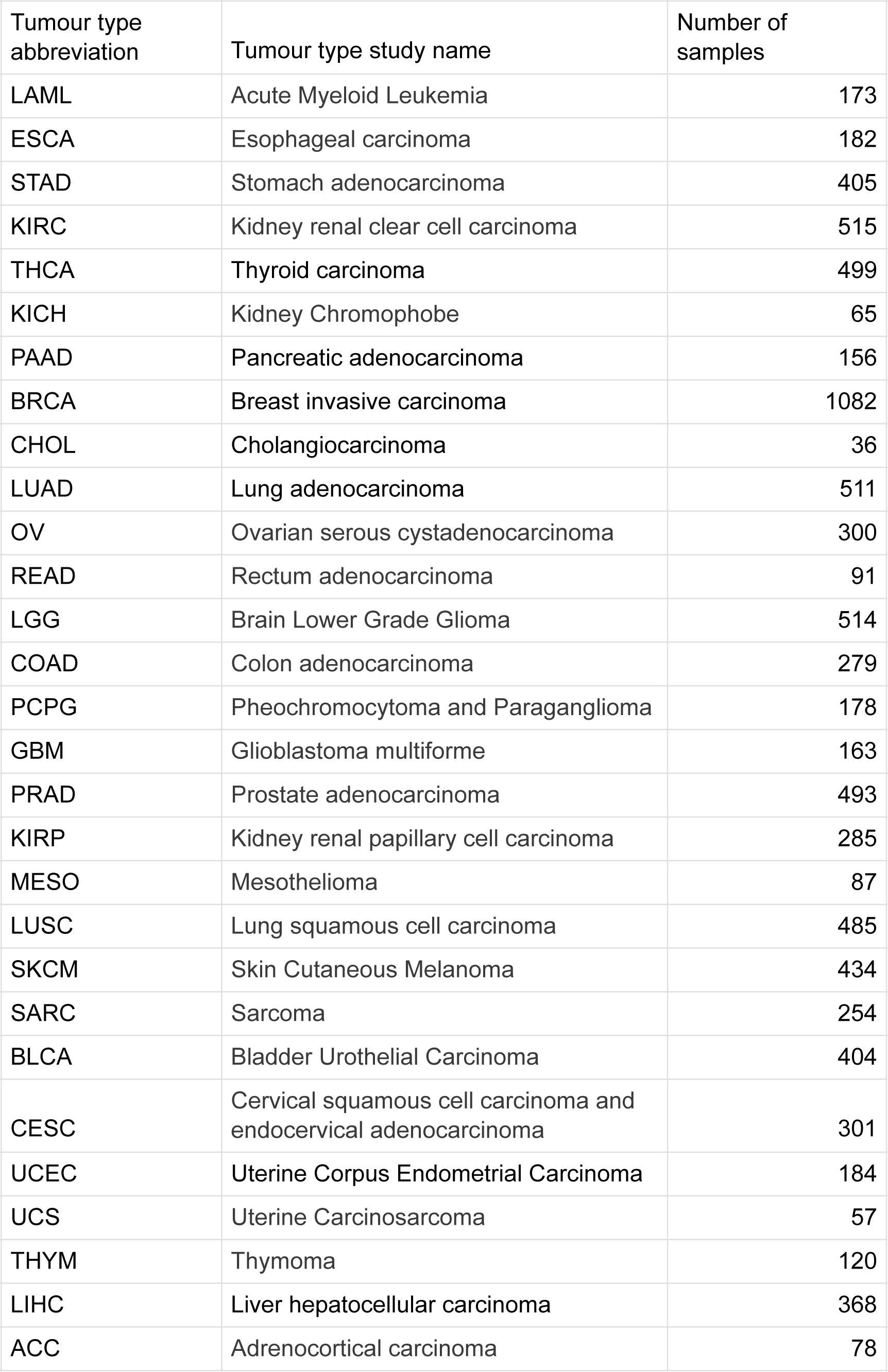
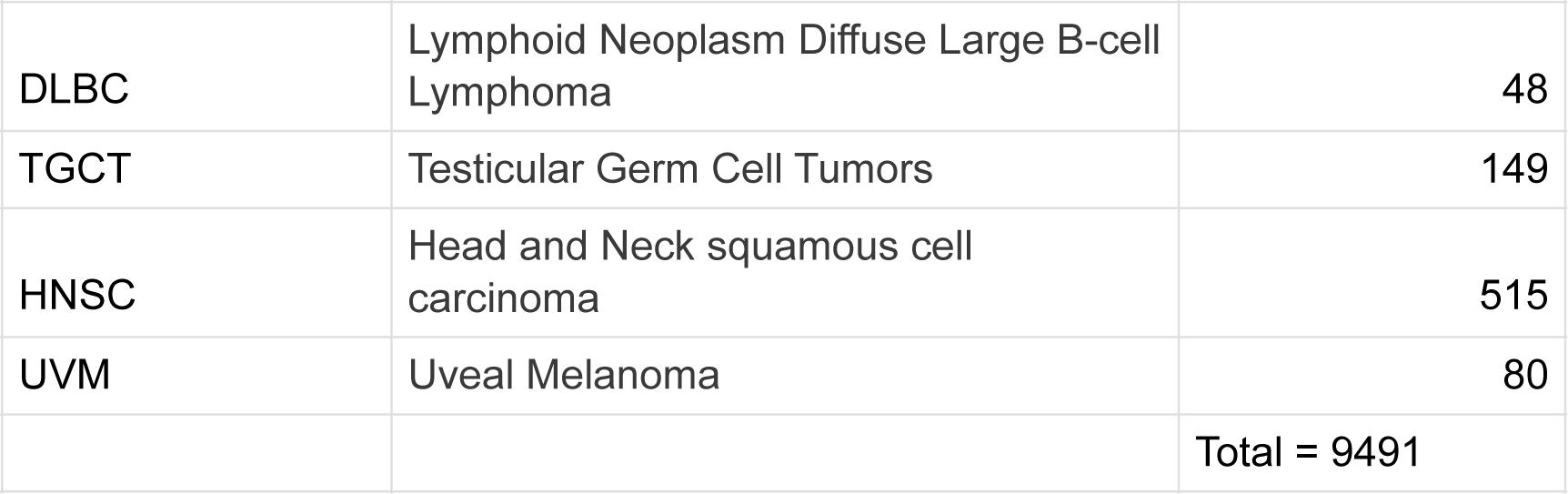
TCGA tumour type abbreviations and full form.

| Tumour type abbreviation | Tumour type study name | Number of samples |
| --- | --- | --- |
| LAML | Acute Myeloid Leukemia | 173 |
| ESCA | Esophageal carcinoma | 182 |
| STAD | Stomach adenocarcinoma | 405 |
| KIRC | Kidney renal clear cell carcinoma | 515 |
| THCA | Thyroid carcinoma | 499 |
| KICH | Kidney Chromophobe | 65 |
| PAAD | Pancreatic adenocarcinoma | 156 |
| BRCA | Breast invasive carcinoma | 1082 |
| CHOL | Cholangiocarcinoma | 36 |
| LUAD | Lung adenocarcinoma | 511 |
| OV | Ovarian serous cystadenocarcinoma | 300 |
| READ | Rectum adenocarcinoma | 91 |
| LGG | Brain Lower Grade Glioma | 514 |
| COAD | Colon adenocarcinoma | 279 |
| PCPG | Pheochromocytoma and Paraganglioma | 178 |
| GBM | Glioblastoma multiforme | 163 |
| PRAD | Prostate adenocarcinoma | 493 |
| KIRP | Kidney renal papillary cell carcinoma | 285 |
| MESO | Mesothelioma | 87 |
| LUSC | Lung squamous cell carcinoma | 485 |
| SKCM | Skin Cutaneous Melanoma | 434 |
| SARC | Sarcoma | 254 |
| BLCA | Bladder Urothelial Carcinoma | 404 |
| CESC | Cervical squamous cell carcinoma and endocervical adenocarcinoma | 301 |
| UCEC | Uterine Corpus Endometrial Carcinoma | 184 |
| UCS | Uterine Carcinosarcoma | 57 |
| THYM | Thymoma | 120 |
| LIHC | Liver hepatocellular carcinoma | 368 |
| ACC | Adrenocortical carcinoma | 78 |
| DLBC | Lymphoid Neoplasm Diffuse Large B-cell Lymphoma | 48 |
| TGCT | Testicular Germ Cell Tumors | 149 |
| HNSC | Head and Neck squamous cell carcinoma | 515 |
| UVM | Uveal Melanoma | 80 |
|  |  | Total = 9491 |

Further, to compare the proteasome expression in a non-malignant state we leveraged 6,125 GTEx samples across 36 tissue types (see Methods). Similar to TCGA tumours, most of the tissue types show significantly (*P*<0.05) higher expression of CP compared to IP (Figure 1B). As expected, the spleen and small intestine had increased IP expression, likely due to the enrichment of lymphocytes in the spleen and inflammation-induced IP expression in epithelial cells in the small intestine (17). However, the positive correlation between CP and IP expression was now only observed in specific tissue types (such as liver, lung, and spleen). On the other hand, the negative correlation was observed in tissue types such as adipose visceral, colon sigmoid, testits and heart-left-ventricle, likely due to the variations in the tissue specific immune environment and physiological conditions (22,23).

Next, to quantify the tumour-driven nature of the proteasomes, we compared the median proteasome expression between tumours (TCGA) and corresponding normal tissues (GTEx) (Figure 1C-E). The IP expression was higher in tumours as compared to matched normal tissues, particularly in cervical (CESC), head-and-neck (HNSC), rectum adenocarcinoma (READ), and bladder adenocarcinoma (BLCA) (Figure 1C). However, this difference was minimal for CP (Figure 1D). Moreover, the median expression difference between the CP and IP was much smaller in tumours compared to the matched normal tissues (Figure 1E), as shown above (Figure 1A, B). Taken together, these results suggest that IP expression is higher in tumour tissues as compared to normal tissues; however, with large variability within and across tumour types.

### 2.2 Immunoproteasome expression is variable among tumour epithelial cells

To delineate the relative contributions of tumour epithelial cells and infiltrating immune cells to the observed CP and IP expression patterns, we explored the publicly available single-cell gene expression data of nine tumour types (see Methods, Supplementary Table 2). As expected, the CP was expressed more in epithelial cells and IP in immune cells (Figure 2A-C, colorectal cancer SMC cohort from (24)). Similar to TCGA tumours (bulk RNA-seq), we observed a significantly (*P*<0.0001) higher CP expression as compared to IP in epithelial cells in all nine cancer types (Figure 2D). Further, in the colon adenocarcinoma (KUL3 cohort from (24)), where tumour-site information was available, we observed that the epithelial cells near the tumour border had higher proteasome expression than cells from the tumour core (median expression fold difference of 1.5 and 2 for CP and IP, respectively) (Figure 2E). This could be due to the higher exposure of tumour cells to the immune cells at the tumour border. In the tumour epithelial population, we also observed a positive correlation between IP and CP expression (Figure 2F, Supplementary Figure 2A), similar to that of bulk transcriptomic data (Figure 1A).

**Figure 2:**
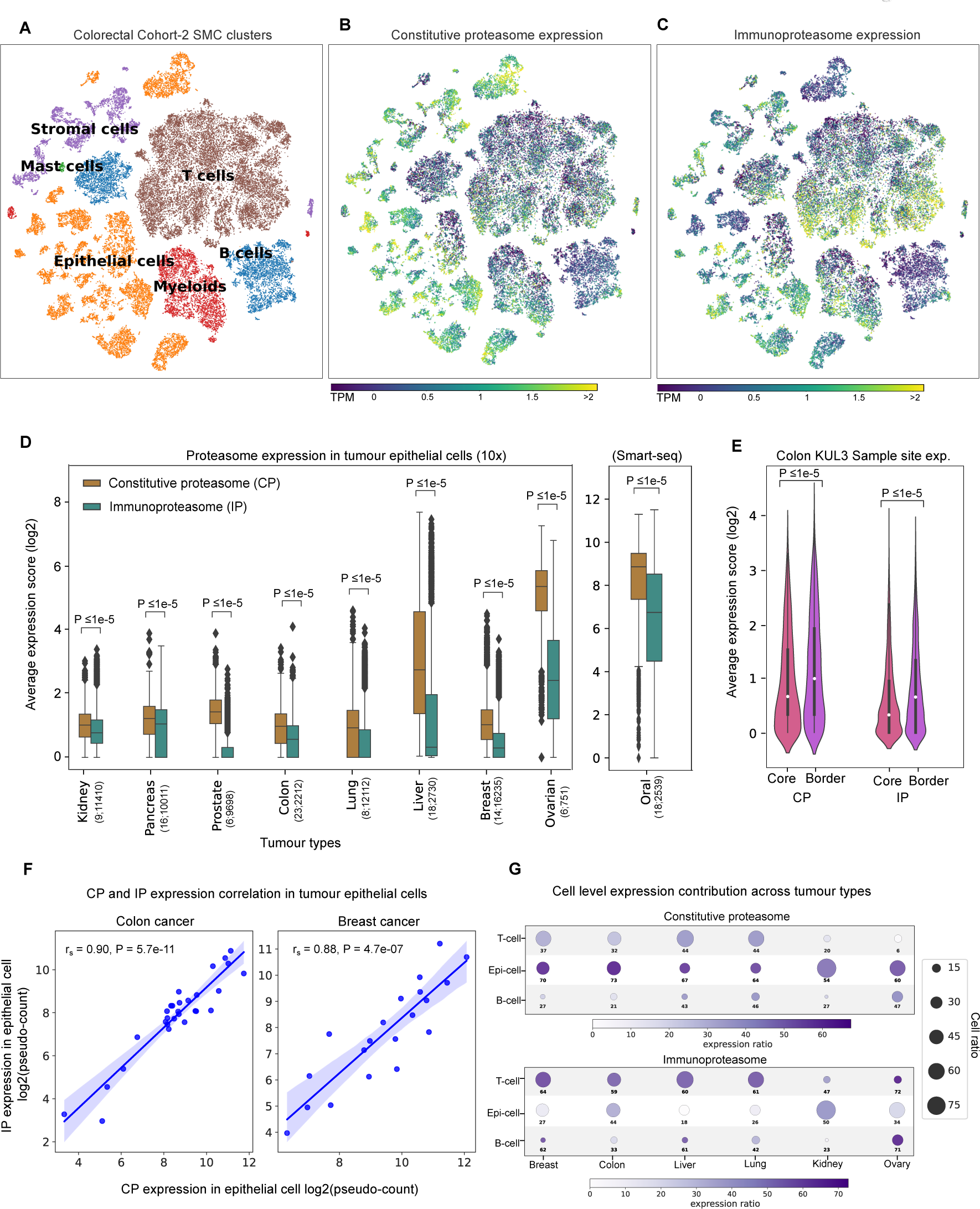
Single-cell analysis of CP and IP expression in tumour epithelial and immune cells. **A.** The t-SNE plot shows the single-cell gene expression data from colorectal cancer. Each dot represents a cell and the colour indicates the cell type annotation (epithelial cells, myeloid cells, B cells, T cells, Mast cells, or stromal cells). **B** and **C** show the average expression of CP and IP, respectively, at the single-cell level. **D.** Distribution of average expression scores of proteasomes (IP and CP) in tumour epithelial cells. The x-axis represents the tumour type and the values within the bracket indicate the number of patients and the number of individual single-cells considered. The y-axis represents the log2 average expression of CP and IP genes. The P-values were calculated using the Mann-Whitney U test (one-sided). **E.** Distribution of average expression scores of CP and IP in tumour epithelial cells with respect to tumour site information – border versus core (represented on x-axis) – in colon adenocarcinoma. The P-values were calculated using the Mann-Whitney U test (one-sided). **F.** Spearman’s rank correlation between the average expression of CP and IP genes (from tumour epithelial cell population) in colon and breast cancers. Each dot represents a sample and the values on the x- and y-axis represent the proteasome expression as pseudo-count (i.e., average expression of CP or IP genes across all cells within the sample). **G.** CP and IP expression level in T-cell, tumour epithelial cell and B-cell population. The size of the circle represents the abundance of the respective cell type in the total cell population under each tumour type. And, the colour inside (and the value mentioned below) represents the percentage of cells having expression value greater than the median expression value in that respective cell type.

Further, we asked what proportion of the tumour epithelial and immune cells express CP and IP. To check this, we selected tumour types annotated with immune cell populations and estimated the proportion of cells expressing proteasome expression above the median level (see Methods) within each group (Figure 2G). This showed that across tumour types, only a subset of tumour epithelial cells (range 18–50%, mean 33%) expressed IP, whereas the majority of the T-cells (range 47–72%, mean 61%) and B-cells (range 23–71%, mean 49%) expressed IP. However, an opposite trend was observed with the CP expression. Taken together, these results suggest that the predominant IP expression is seen only in a subset of the tumour cells (∼33%). Also, the expression level varies among the tumour cells with respect to their spatial distribution (tumour border vs core).

### 2.3 Tumours with high immunoproteasome expression are enriched with anti-tumorigenic immune cells

To study the impact of tumour-infiltrating immune cells (TIL) on the expression of IP, we performed a differential analysis of enrichment of 18 different types of immune cells between high-IP (top quartile) and low-IP (bottom quartile) expression sample groups under each tumour type, by using GSVA (see Methods, Supplementary Table 3). With the GSVA enrichment score difference of > |0.2| and FDR adjusted p-value < 0.1, we found the enrichment of activated CD8+ T cells, activated dendritic cells (aDC), exhausted T cells and cytotoxic cells (which represents the anti-tumour activity of overall immune cell infiltrates, see Methods) in high-IP groups across multiple cancer types (Figure 3A). These immune cells are known to produce inflammatory cytokines (IFN-γ and TNF-α) that can induce IP expression in tumour cells. The enrichment of exhausted T cells in high-IP groups could be explained by persistent antigen presentation and chronic inflammation in the tumour microenvironment (TME) (25). In line with this, we observed a positive enrichment of activated dendritic cells (aDC) compared to immature dendritic cells (iDC) in multiple tumour types (Figure 3A). But, the positive enrichment of regulatory T cells (Tregs) observed in certain tumour types could be explained by the fact that Tregs creates an immunosuppressive environment and favour tumour cells for immune evasion (26). Also, the induction of IP in immune cells (such as dendritic cells and T cells) plays a key role in the dendritic cell activation process (27,28) and T cell differentiation (14,29). On the other hand, the eosinophils and T-helper cells were negatively enriched in the high-IP-expression group.

**Figure 3:**
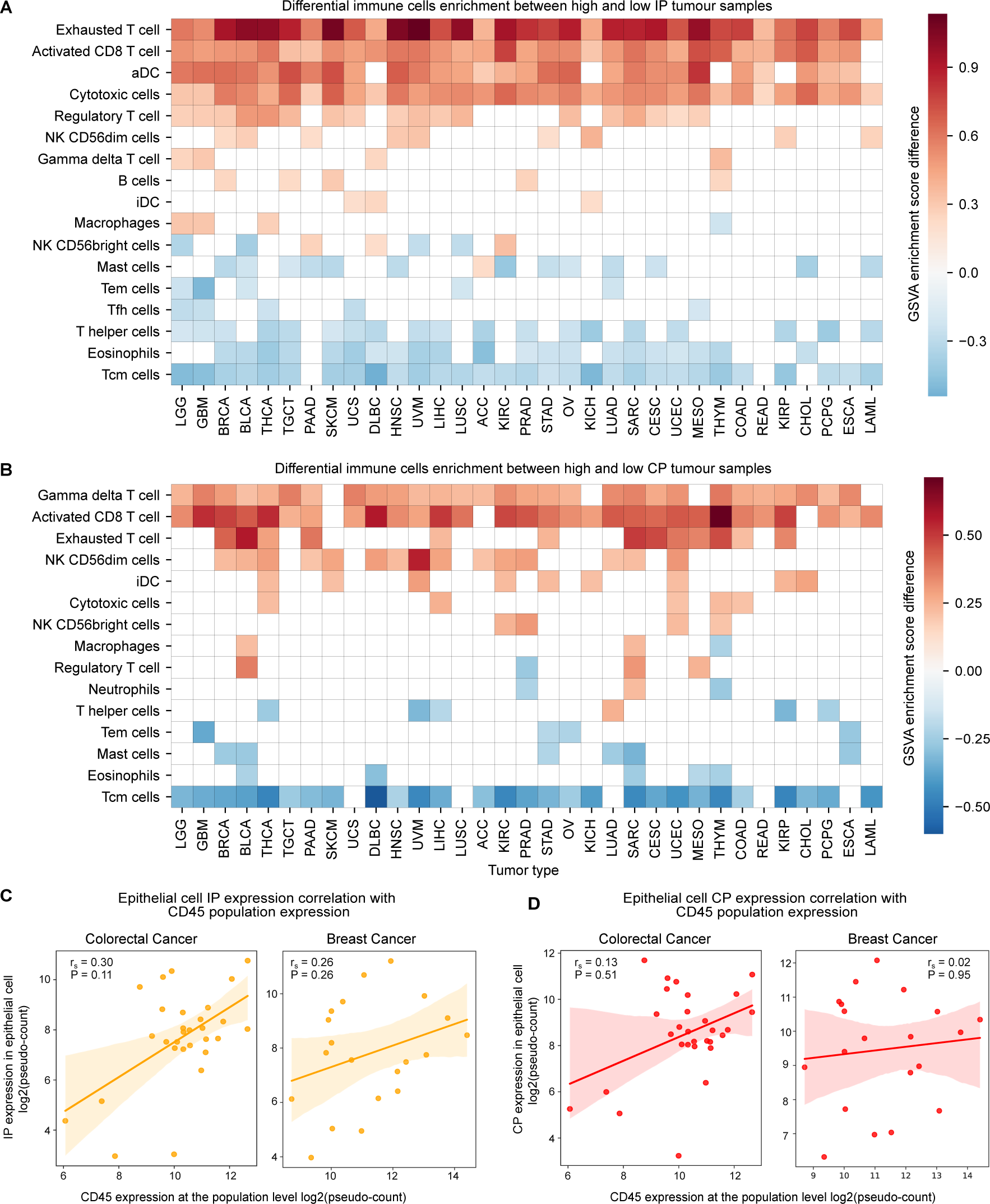
Immune cells associated with CP and IP expression. **A.** Heatmap represents the differential enrichment of immune cell types between sample groups of high-IP (upper quartile) and low-IP (bottom quartile) expression under each tumour type (see Methods). The x-axis represents the tumour type and the y-axis represents the different immune cell types. The colour of each cell represents the GSVA enrichment score difference between the high-IP versus low-IP expression group. Only those immune cell types that showed significant differences (GSVA score difference of >|0.2| and FDR adjusted p-value < 0.1) and appear in more than two tumour types were plotted here. The cells with white colour indicate no significant enrichment with respect to the above condition. **B.** Same as **A**, but for the CP. **C.** Spearman’s rank correlation between CD45 gene expression (immune cell marker) among all cells within the tumour and the average IP expression in epithelial cells at the sample level in colorectal and breast cancers (by using single-cell gene expression data). **D.** Same as **C**, but for the CP.

In the case of CP, the gamma-delta T cells showed positive enrichment (Figure 3B) across tumour types, likely because of the extensive repertoire of these cells within the epithelial tissues (30). Similar to the IP, the activated CD8+ T cells were positively enriched in the high-CP group in multiple tumour types (Figure 3B), consistent with the expression correlation observed between CP and IP in bulk (Figure 1A) and single-cell gene expression analysis (Figure 2F). The central memory T (T_CM_) cells were negatively enriched in both the CP and IP groups.

Further, to investigate the effect of immune cells on the expression of proteasome in the tumour epithelial cells at the single-cell level, we compared proteasome expression with CD45 expression (see Methods), a transmembrane glycoprotein expressed in all lymphocytes (31). The IP expression from tumour epithelial cells was highly correlated with CD45 expression from whole-cell populations in colorectal, breast, lung, and oral cancer (Figure 3C, Supplementary Figure 2B). This can be attributed to the enrichment of cytotoxic immune cell infiltrations (as seen above); thus with an increase in the lymphocyte population we can see an increase in IP expression also. There was also a positive CP correlation with the increase in the lymphocyte population, but the association strength was much weaker as compared to IP (Figure 3D, Supplementary Figure 2C). However, in kidney (renal cell) cancer, a negative correlation was observed for both IP and CP expression with the CD45 expression (Supplementary Figure 2B,C), which could be due to the lack of cytotoxic or interferon signalling in the TME.

Taken together, these results suggest that the presence of specific immune cells with cytotoxic activity correlates positively with IP expression in tumour epithelial cells. This can potentially enhance antigen presentation in tumour cells and attract more cytotoxic immune cells, like a positive feedback mechanism.

### 2.4 High immunoproteasome expression is associated with inflammatory response and oxidative stress

Following the characterisation of TILs, we further asked which pathways were associated with the IP expression. For this, we performed differential pathway analysis of 50 MsigDb hallmark gene-sets between high-IP (top quartile) and low-IP (bottom quartile) sample groups under each tumour type, by using GSVA (see Methods, Supplementary Table 4). With the GSVA enrichment score difference of > | 0.2| and FDR adjusted p-value < 0.01, we identified seven upregulated and six downregulated pathways, recurrent in more than 40% of the tumour types (Figure 4A). The most frequently upregulated pathways are interferon-alpha (in 32 cancer types) and interferon-gamma (in 31 cancer types). Interferon signalling is known to induce IP formation in both immune and non-immune cells. Among all tumour types, DLBC and THYM did not show a significant increase in interferon-alpha and/or -gamma signalling, which could be attributed to the high basal level expression of the IP in these tumour types. The other pathways of upregulation in multiple cancer types are allograft rejection (*n* = 28), IL-6/JAK/STAT3 signalling (*n* = 24), inflammatory response (*n* = 23), ROS pathway (*n* = 22), and TNF-α signalling through NF-kB (*n* = 14). The allograft rejection pathway has several overlapping genes with interferon signalling, which could explain their frequent upregulation. Upregulation of the ROS pathway with high IP expression is likely due to oxidative damage within the tumour cells and enrichment of inflammatory cells in the TME (32). TNF-α is a pro-inflammatory cytokine previously shown to induce the expression of IP subunits, either in synergy with IFN-γ or independently in a tissue-specific manner (33). But, none of these pathways (except ROS) were enriched in high-CP tumour samples as compared to low-CP samples (Supplementary Figure 3A).

**Figure 4:**
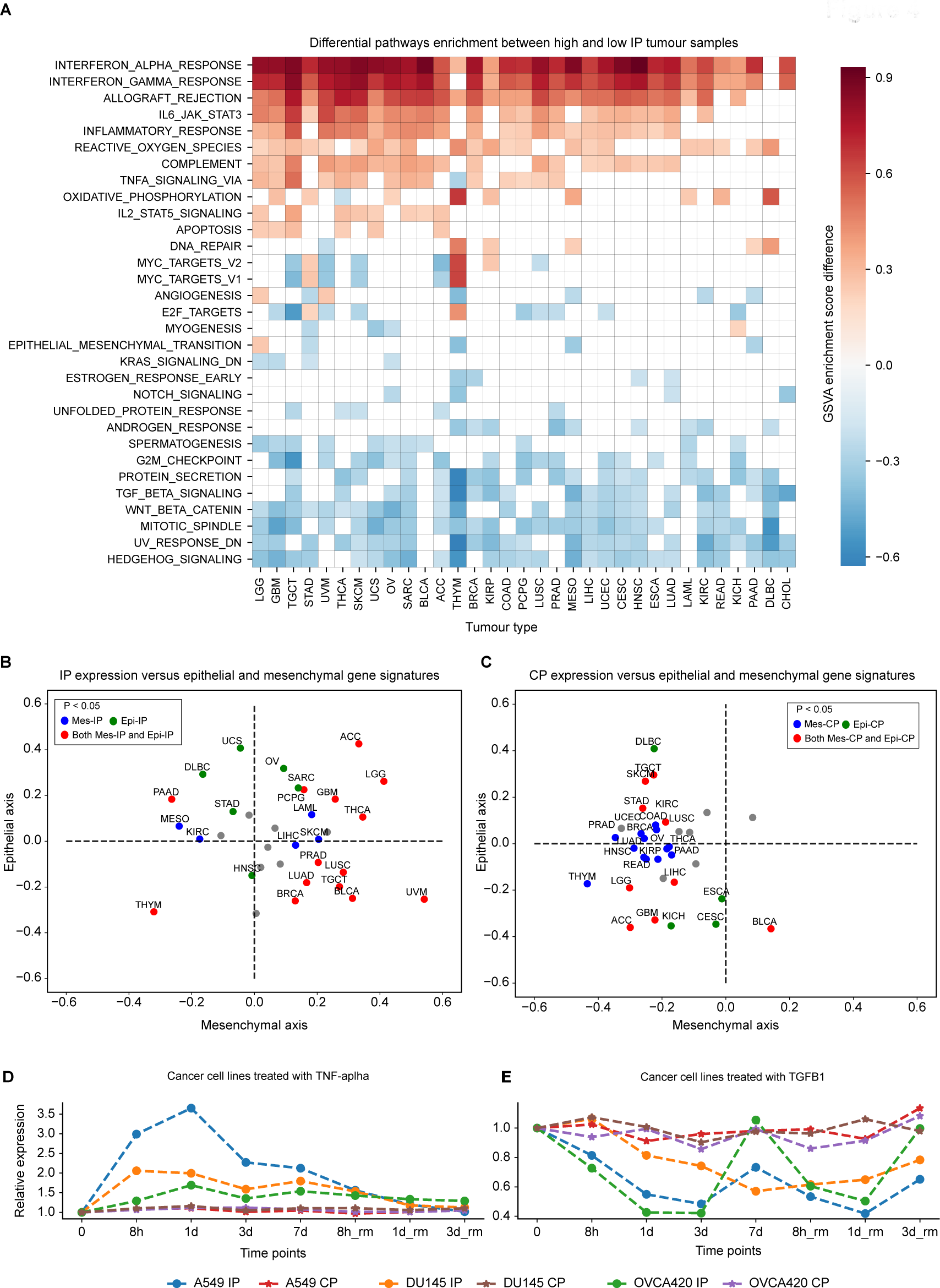
Pathways associated with proteasome expression. **A.** Differential enrichment of hallmark gene sets (from MsigDB) between sample groups of high-IP (upper quartile) and low-IP (lower quartile) expression under each tumour type. The x-axis represents the tumour type and the y-axis represents the different hallmark pathways. The colour of each cell represents the GSVA enrichment score difference between the high and low IP expression group. Only those pathways that showed significant differences (GSVA score difference of >|0.2| and FDR adjusted p-value < 0.01) and appear in more than three tumour types were plotted here. The cells with white colour indicate no significant enrichment with respect to the above condition. **B.** Correlation of average IP expression with the epithelial specific (x-axis) and mesenchymal (y-axis) specific gene signatures. Each dot is a tumour type and the colour indicates whether the correlation with either the epithelial specific or mesenchymal specific gene signature, or both are significant (P<0.05) or non-significant (grey colour). **C**. Same as **B**, but for the CP. **D-E.** Relative expression changes in the IP and CP expression level in three different cancer cell lines (A549, lung; DU145, prostate; OVCA420, ovarian) treated with TNF-α (**D**) and TGF-β1 (**E**), respectively. The x-axis represents the different time points: untreated (0), under treatment (8h, 1 day, 3 days, 7 days) and withdrawal (rm) of treatment after 7 days of treatment (8h, 1 day and 3 days). The y-axis represents the relative expression change with respect to untreated condition (0). Each cancer cell line and proteasome (IP and CP) combination is highlighted with different lines.

Among the downregulated pathways with IP expression, the hedgehog (Hh) signalling (*n* = 26), mitotic spindle (*n* = 25), UV response (*n* = 25), Wnt-beta signalling (*n* = 23), TGF-β signalling (*n* = 20), and G2M checkpoint genes (*n* = 17) were most frequent between the different cancer types (Figure 4A). The association of Hh signalling with low IP expression could be due to the suppression of CD8+ T cell infiltration in the TME through the sonic hedgehog (Shh) mediated polarisation of tumour-associated macrophages (TAMs) (34). Similarly, the activation of Wnt-beta signalling has been shown to prevent the neoantigen-specific CD8+ T infiltration into the TME in NSCLC (35). TGF-β functions as an anti-inflammatory cytokine and it has been shown to downregulate the IP expression in NSCLC (15). Together, these results suggest that the IP expression in tumours is negatively correlated with the immunosuppressive environment, which could be triggered by the tumour cell-intrinsic pathways and the types of immune cell infiltrating the TME.

A previous study has shown that the epithelial-to-mesenchymal transition (EMT) negatively correlated with IP expression in NSCLC (15). However, our analysis found that the EMT pathway was downregulated in only four cancer types (Figure 4A). To further investigate this, we compared the expression of CP and IP with the epithelial-specific and mesenchymal-specific gene signatures (see Methods). The correlation of IP expression with mesenchymal signature was more distributed (Figure 4B): negative in certain tumours (HNSC, KIRC, PAAD, STAD, and UCS) and positive in others (PRAD, GBM, THCA, LUSC, PCPG, and BLCA). But, the CP expression was negatively correlated with mesenchymal-specific signatures (Figure 4C) and EMT (Supplementary Figure 3A) in multiple cancer types. Together, these results suggest that EMT, particularly the upregulation of mesenchymal signatures, downregulates the IP expression in a tumour-type specific manner as compared to CP.

### 2.5 Immunoproteasome expression is strongly associated with TNF-α and TGF-β signalling

From the above analysis, we see certain pathways are consistent and strongly associated with up- and downregulation of IP expression. However, it does not reveal how long it would take for these signalling cascades to induce or suppress the IP expression. To study this, we used the available time-course data of four different cancer cell lines (A549, lung; DU145, prostate; OVCA420, ovarian; MCF-7, breast) treated with TNF-α (positive regulatory of IP) and TGF-β1 (negative regulatory of IP) (36). We observed that the treatment with TNF-α led to an increase in IP expression, with a roughly 2- to 3-fold increase at 8 hours or one-day post treatment, but not at the cost of CP expression level going down (Figure 4D). On the other hand, TGF-β1 treatment led to a decrease in the expression, roughly 2-fold on day one or three days post-treatment, also without any difference in CP expression. This suggests that the induction and suppression rates of IP expression differ between TNF-α and TGF-β1, respectively. Nevertheless, in both cases, removal of cytokines brought the proteasome expression level closer to its pre-treatment level (with some differences between cell lines, particularly for TGF-β1). An exception to this is the MCF-7 cell line (Supplementary Figure 3B), which showed increased IP expression even after removal of TNF-α at 8 hours, and increased IP expression with TGF-β1 treatment (in contrast to other cancer cell lines as shown above).

### 2.6 Immunoproteasome gene expression is not correlated with somatic copy number changes

Next, we asked whether the proteasome (CP and IP genes) were targeted by somatic genetic alterations (mutations and/or copy number alterations) that favour the tumour cells for immune evasion. Surprisingly, we found that only a few samples (< 1%) harbour protein-affecting mutations (after removing hypermutator samples see Methods, Supplementary Table 5). However, low-level somatic copy number alterations were seen across different tumours (Supplementary Figure 4A). A significant fraction of samples (above 50%) showed low-level amplification and deletions in both CP and IP across tumour types (KICH, UCS, OV, and BRCA). Deep deletion was observed in DLBC, albeit at a lower proportion of samples (∼10%). Further, we asked whether the copy number alterations also lead to changes in gene expression. As expected, we observed a strong correlation between copy number changes and expression for each CP gene. However, for IP genes, we did not observe a strong association at the pan-cancer level (Supplementary Figure 4B) nor at the individual tumour type level (Supplementary Figure 4C). To understand this further, we compared the expression level of upstream regulators (IL-6/JAK/STAT3 and interferon-gamma pathways) of IP genes between samples with and without copy number alterations of IP genes. This showed that overall expression of the above regulators was significantly lower in the samples with gene copy amplification, as compared to samples without alterations (Supplementary Figure 4D). This could be due to the hypermethylation or genetic alterations in the upstream regulatory genes. Together, these results suggest that the expression of IP genes is dependent on the activity of the upstream regulators and not the gene copy number alone.

### 2.7 Association of immunoproteasome expression with overall survival is influenced by the immediate immune environment

We asked how IP expression influences patient survival using the TCGA cohort. Based on the Cox regression analysis, we found that the higher expression of IP was associated with better overall survival (FDR adjusted *P*<0.1) in six tumour types: skin cutaneous melanoma (SKCM), cervical squamous cell carcinoma (CESC), breast invasive carcinoma (BRCA), bladder urothelial carcinoma (BLCA), mesothelioma (MESO), and sarcoma (SARC). On the other hand, the expression of IP was associated with poor survival in kidney renal clear cell carcinoma (KIRC), acute myeloid leukaemia (LAML), brain lower grade glioma (LGG), and uveal melanoma (UVM) (Figure 5A).

**Figure 5:**
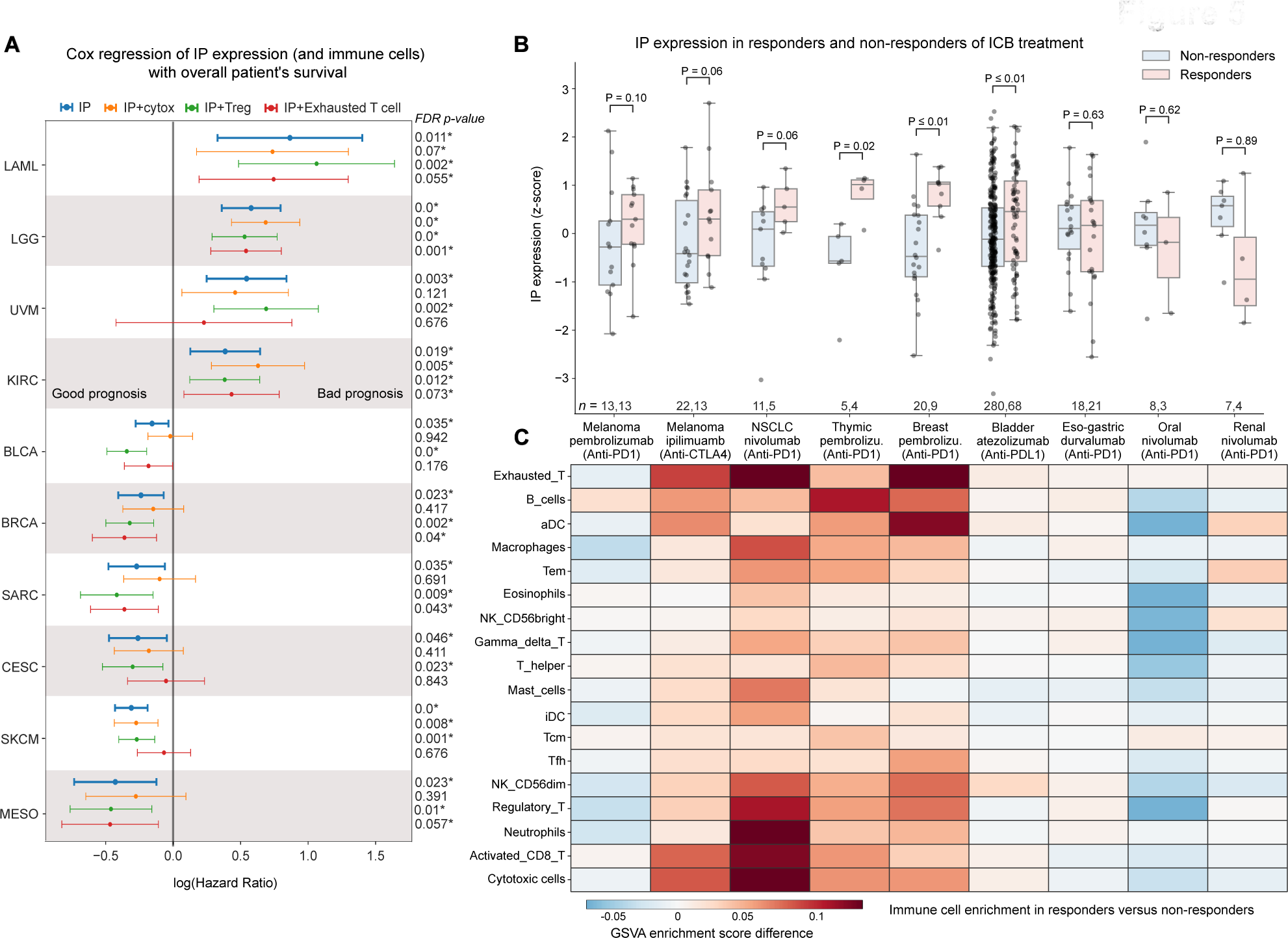
Association of IP expression with overall survival and response to immune checkpoint blockade (ICB) therapies. **A.** Cox proportional hazard ratio (HR) for IP expression alone and combined with three other confounders (cytotoxic cells, regulatory T cell and exhausted T cells) in 10 different tumour tissues from TCGA which showed significant association. The x-axis represents the log(HR) value (> 0 indicates bad prognosis and < 0 indicates good prognosis), and the horizontal line represents the mean and 95% confidence interval. The colour of the line indicates the variables considered for the analysis. The FDR adjusted p-value shown on the right was calculated using the Wald test and corrected for multiple hypothesis testing using Benjamini-Hochberg method. The asterisks(*) symbol indicates FDR adjusted p-value less than 10% significance level. **B.** The boxplot shows the average IP expression (represented as z-scores) in responders versus non-responders of ICB therapies in nine different tumour types. Each data point in the box plot represents a sample, the horizontal middle line indicates the median, the height of the shaded box indicates the interquartile range (IQR), and the whiskers indicate 1.5 x IQR. The P-values shown at the top were computed using a one-sided Mann-Whitney U test (testing whether IP expression level in responder was higher than non-responder or not). **C.** Heatmap showing the differential enrichment score difference of 18 different immune cells (based on its gene signature score) between responders and non-responders of ICB therapy.

Given that the IP expression in bulk RNA-seq analysis has contributions from the infiltrating immune cells, we tried to adjust for their effects and other confounders such as age, gender, tumour stage, and tumour mutational burden (TMB) of the samples independently and combined using multivariate Cox regression analysis (see Supplementary Table 6). This showed that the enrichment of different types of immune cells strongly influences the association of IP with survival in the above tumour types (as compared to other confounders). These immune cell types can be broadly classified into having anti- or pro-tumorigenic potential based on their functional properties. The anti-tumorigenic group includes cytotoxic cells such as CD8+ T cells, gamma-delta T cells, and natural killer cells, whereas the pro-tumorigenic group includes regulatory T cells (Tregs) and exhausted T cells. When we adjusted for the enrichment of anti-tumorigenic (cytotoxic) immune cells, the hazard ratio (HR) increased and became non-significant (except for melanoma). This suggests the overwhelming role of cytotoxic immune cells (along with induction of IP expression in tumour cells) towards better overall survival rates of patients.

On the other hand, with adjustment for the effect of Tregs, which are immunosuppressive in nature, the association of IP with better survival improved for the above tumour types, as well as for LGG, which showed a bad prognosis with IP expression alone (Figure 5A). In the case of melanoma, the adjustment for neither cytotoxic cells nor Tregs impacted the association of IP with better survival, suggesting a cell-intrinsic or inherent higher expression of IP in this tumour type that favours better survival (19). Whereas, when we adjusted for exhausted T cells (which have poor effector/tumour-eliminating function), the tumour types that showed poor survival with IP expression improved, and those tumour types with good survival (including melanoma) became worse, suggesting the presence of exhausted T cell plays a different role in a tumour-type specific manner. With CP expression, we did not find any tumour type (except LGG) that showed good prognosis (Supplementary Figure 5A). Together, these results suggest that IP expression is better associated with the overall survival of patients, however, it is strongly influenced by the infiltrating immune cell types and their activity towards pro- or anti-tumorigenicity.

### 2.8 Association of immunoproteasome expression with response to immune checkpoint blockade therapies is tumour-type specific

Next, we asked if IP expression can be used as a predictor for immune checkpoint blockade (ICB) treatment response. To test this, we used data from nine publically available studies across eight tumour types (melanoma, lung, thymus, eso-gastric, breast, bladder, oral and renal) (see Methods, Supplementary Table 7). We compared the average IP expression (z-score) between the responders and non-responders on treatment with ICB (Figure 5B, Supplementary Table 8). In skin cutaneous melanoma, for both anti-PD-1 (Pembrolizumab) and anti-CTLA-4 (Ipilimumab), we see responders having higher IP expression pre-treatment. This could be due to the enrichment of activated CD8+ T cells and cytotoxic cells in responders as compared to non-responders (Figure 5C) (37) (19). Similar trends can be seen in non-small-cell lung cancer (NSCLC) (Figure 5B), where patients with high IP expression respond better to the anti-PD-1 treatment (Nivolumab). They also show high enrichment of neutrophils, activated CD8+ T cells, and Tregs as compared to non-responders (Figure 5C). This enrichment profile matches with the hot immunophenotype of NSCLC cancer defined by Lizotte et al (38). A significant difference in the IP expression between responders and non-responders can be seen in thymic carcinoma (Figure 5B). Although the thymic epithelial tumours have the lowest TMB among adult cancers (39), the dysfunction in the T-cell development creates an autoreactive T cell population. These autoreactive T cells can recognise self-antigens and are known to increase IFN-γ signalling, which leads to the induction of IP genes and checkpoint receptors that can be beneficial for ICB response (40). Similarly, breast cancer patients (Figure 5B) also show high IP expression in responders as compared to non-responders of anti-PD-1 therapy (Pembrolizumab). This is in line with the previous study (41) that showed enrichment of IFN-γ, antigen presentation and checkpoint receptor gene sets in expanding T cells with effector function on ICB treatment. Also, in metastatic bladder urothelial cancer, the patients who responded well for anti-PDL-1 (atezolizumab) showed high IP expression as compared to the non-responders. This could be explained by the enrichment of CD8+ T cells and high tumour mutation burden in these tumours (42). Moreover, we see an overall enrichment of exhausted T cells (43) (41) in cancer types that have a positive association of IP expression with ICB response (Figure 5C). This suggests that some of these exhausted T cells might reactivate and gain effector function upon treatment (44).

However, we observed variability in the IP expression association with response to ICB therapy in other solid tumour types. In eso-gastric cancer, we did not see a difference in the IP expression and immune cell enrichment between the responders and non-responders of anti-PD-1 treatment (Figure 5B). In oral and renal cancers, we observed lower IP expression in responders as compared to non-responders of anti-PD-1 (Nivolumab) treatment. In the case of oral cancer, the downregulation of IP could be due to the enrichment of Tregs in the non-responders (Figure 5C). In contrast, in renal cancers, we observed a relatively higher IP expression in non-responders, despite their poor response to anti-PD-1 therapy. This could be due to the low enrichment of cytotoxic immune cells in the TME (45) or other tumour-cell intrinsic factors (46). With CP expression, we did not find any significant difference between responder and non-responder across tumour types (Supplementary Figure 5B). Taken together, these results suggest that IP expression is strongly associated with response to ICB; however, in a tumour-type specific manner that can be due to the enrichment of tissue- and tumour-type specific immune infiltration patterns and their (tumour-eliminating) effector function. Thus, IP expression combined with immune infiltration patterns could serve as a better predictor for ICB response in solid tumours.

## 3. Discussion

Though immune checkpoint blockade (ICB) therapies are emerging anti-cancer therapy, the response rate is still variable within and across solid tumour types. To gain insight into this, recent studies have focused on the characterisation of genetic and molecular alterations at the tumour and immune-cell levels, and whether these alterations can be used as biomarkers to predict response to ICB therapies (45,47,48). For example, the study by Kalaora et al. (19), showed that IP gene expression can be used as a potential biomarker to predict prognosis and response to ICB therapy in skin melanoma. However, the extent of this in other solid tumours remains unclear. Thus, in this study, we explored the expression patterns of IP (in relation to CP) across multiple solid tumour types and identified cell-extrinsic and cell-intrinsic factors that influence the induction of IP expression and their association with the patient’s overall survival and response to ICB therapies.

Although previous works have shown that the IP expression can be induced in non-immune cells upon exposure to pro-inflammatory signalling or stress conditions (1,6), it is not clear whether the induction of IP can affect the expression of CP. Here, we show that the expression of IP and CP were positively correlated in multiple tumour types (as compared to normal tissue) (Figure 1A, 2F); however, the induction level of IP was higher than the CP (Figure 4D, E). We speculate that this could be due to potential co-regulation or cross-talk between the CP and IP gene expression in solid tumours (as previously shown in myeloma (49)). In addition, this observation raises the question of whether tumour cells require both CP and IP (or intermediate form) to modulate the antigen presentation depending on the stress from infiltrating immune cells. Future studies involving proteomics combined with epitope mapping could help to answer these questions and to understand the dynamics of active CP and IP complexes in shaping the antigen presentation and immunogenicity in tumour cells.

In most of the solid tumour types, IP expression was positively correlated with cytotoxic immune cell infiltration and upregulation of genes involved in interferon signalling pathways and reactive oxygen species (Figure 3 and 4), suggesting that the induction of IP is influenced by both cell-extrinsic and cell-intrinsic pathways. However, within the tumour tissue, the expression of IP was highly variable among tumour epithelial cells: only a subset of cells expressing high IP, in particular, those at the tumour border rather than tumour core regions (Figure 2 E and G). Recent studies have shown that the spatial enrichment of cytotoxic immune cells plays a vital role in interferon signalling and response to ICB therapies (47,50). Based on this, we speculate that the variation observed in IP expression among tumour cells may be due to the differential exposure of tumour cells to the cytotoxic immune cells and interferon signalling. However, future studies involving spatial-temporal analysis of immune-cell infiltration patterns and cell–cell interactions across different cancer types could reveal the complex interaction between the tumour cells and immune cells, and their impact on induction of IP expression.

Finally, we showed that the association of IP expression with overall survival rate and response to ICB therapies is tumour-type specific and is greatly influenced by the tumour infiltrating immune cells (Figure 5). This result can help to identify the tumour types suitable for ICB therapies or in combination with proteasome inhibitors (or other targeted therapies) to further enhance the clinical response. For instance, in tumour types (such as skin melanoma, breast, NSCLC, bladder and thymus) where IP expression was associated with better prognosis and response to ICB therapies, immunoproteasome activators (IFN-γ or 5-aza-dC) or others to enhance interferon signalling can be combined with ICB therapies (15). In particular, for those patients with low or moderate IP expression in these tumour groups. On the other hand, the tumour types (such as glioma, acute myeloid leukaemia, oral and renal) where IP expression was associated with poor prognosis and non-response to ICB therapies, treatment with immunoproteasome inhibitors, or chemotherapy along with ICB can be explored to enhance immune response (18,51,52). Although the proteasome inhibitor has shown promising results in treating haematological cancers, its potential for treating solid cancers requires further explorations. For example, treatment of glioblastoma cell line with ONX-0914, a selective immunoproteasome inhibitor targeting PSMB8, has shown potential to reduce tumour progression by inducing cell cycle arrest and autophagy (53). Alternatively, the development of CAR T cells in combination with oncolytic viruses can help to target the immunosuppressive tumours to enhance antigen presentation and cytotoxic T cell response (54).

In sum, our study reveals that the expression of IP, combined with immune cell infiltration patterns, can be used as a potential marker to predict prognosis and response to ICB therapies in solid tumours. However, further studies with more samples and cancer types with response data of ICB therapies are required to validate these findings and also to explore whether the IP expression has any oncogenic roles in these tumours besides their expected role in protein degradation.

## 4. Methods

### 4.1 TCGA and GTEx data processing

Level 3 pre-processed gene expression data (illuminahiseq_rnaseqv2, RSEM_genes_normalized) of 10,446 TCGA samples (from 9,695 unique patients across 33 different tumour types) were downloaded from the Broad GDAC firebrowse (http://firebrowse.org/, TCGA data version 2016_01_28). The samples flagged for quality control issues (“Do_not_use” == True, according to “Merged Sample Quality Annotations” recommendations, https://gdc.cancer.gov/about-data/publications/pancanatlas) were filtered out. For patients with multiple samples, only one sample (preferably sample type code 01 – primary tumour) was considered. In skin cutaneous melanoma (SKCM), the samples from distant metastases were excluded to focus on primary tumours and regional metastasis (55). The final list of 9,491 unique patient tumour samples included in this study were given in the Supplementary Table 1. For these samples, the corresponding somatic mutations (MC3, “mc3.v0.2.8.PUBLIC.maf.gz”), copy-number alterations (GSTIC2.0, “all_thresholded.by_genes_whitelisted.tsv”) and tumour purity (ABSOLUTE, “TCGA_mastercalls.abs_tables_JSedit.fixed.txt”) information were obtained from the TCGA pan-cancer atlas study (https://gdc.cancer.gov/about-data/publications/ pancanatlas, https://gdc.cancer.gov/about-data/publications/pancan-aneuploidy). For GTEx, the pre-processed gene expression data (RSEM gene normalised, which were processed and normalised similar to TCGA), of 6,125 normal tissues (across 36 tissue types) was downloaded from the UCSC Xena browser (https://toil-xena-hub.s3.us-east-1.amazonaws.com/download/gtex_RSEM_Hugo_norm_count.gz)

(56). The expression (RSEM) values were log2 transformed (after adding 1) and then computed as an average expression value for CP (mean of PSMB5, PSMB6, and PSMB7 genes) and IP (mean of PSMB8, PSMB9, and PSMB10 genes) at each individual sample level (Figure 1A,B, Supplementary Table 1). The relationship between the average expression of CP and IP was estimated using Spearman’s rank correlation approach. The matched normal tissue (from GTEx) for each tumour type shown in Figure 1C-E was selected based on the information given in the Supplementary Table 1 of Tamborero et al. (57).

### 4.2 Single-cell data processing

We have collated single-cell gene expression data from previous studies covering nine tumour types: kidney (*n* = 9), pancreas (*n* = 16), prostate (*n* = 6), colorectal (SMC cohort, *n* = 33; KUL3 cohort, *n* = 18), lung (*n* = 17), breast (*n* = 14), liver (*n* = 18), ovarian (*n* = 6) and oral (*n* = 18). The number of sample types (tumour and adjacent normal), single-cell counts, data format, and the study source were given in Supplementary Table 2. For studies that provided raw UMI counts, the counts were normalised at cell level by the total counts over all genes using scanpy (58) (sc.pp.normalize_total function with target_sum = 1e4). Further, the values were log2 transformed (after adding 1) and then computed as the average expression value for CP (mean of PSMB5, PSMB6, and PSMB7 genes) and IP (mean of PSMB8, PSMB9, and PSMB10) at each cell. To calculate the proportion of cells expressing IP and CP under different cell types (tumour epithelial cells, T cells, and B cells) shown in Figure 2G, we selected tumour types that have those three cell types available and calculated the fraction of cells with average proteasome expression (IP or CP) greater than the median average proteasome expression value (IP or CP) in the respective cell types. Similarly, for the sample-level comparison (shown in Figures 2F and 3C), we considered cells with average proteasome expression values above the median value in the respective cell type and calculated the sum of the average proteasome expression values across all cells within that sample (represented as pseudo-counts). Spearman’s rank correlation analysis was used to compare proteasome expression at the sample level in Figures 2F and 3C.

### 4.3 Enrichment analysis of immune cells and pathways

We performed a single-sample gene set enrichment analysis (ssGSVA) using R package GSVA (59) to quantify the infiltration level of 18 different immune cells (cytotoxic cells, regulatory T cells, gamma-delta T cells, activated CD8+ T cells, iDC, aDC, Tfh cells, T_em_ Cells, T_cm_ Cells, T-helper cells, neutrophils, NK CD56^dim^ cells, NK CD56^bright^ cells, mast cells, macrophages, eosinophils, B cells, and exhausted T cells) by using their respective gene signatures from Tamborero et al., (57), except for the exhausted T cell for which we considered three genes (LAG3, HAVCR2, and PDCD1) from (41,60). The cytotoxic cells is a meta group that represents the anti-tumour activity of overall immune cell infiltrates by considering highly expressed genes from activated CD8+ T cells, gamma delta T cells and natural killer (NK) cells (57). The 50 different hallmark pathways were obtained from MsigDB (61). The differential pathways and immune cell enrichment between high (> 75th percentile) and low (< 25th percentile) proteasome of IP or CP groups were carried out through the GSVA pipeline (59). The GSVA enrichment score difference of >|0.2| and FDR adjusted *p*-value < 0.1 was considered significant. The gene signatures associated with epithelial-specific and mesenchymal-specific were obtained from Tan et al. (62). These gene lists were given in Supplementary Table 9.

### 4.4 Cancer cell lines data analysis

The time-course data of four cancer cell lines (A549, lung; DU145, prostate; OVCA420, ovarian; and MCF-7, breast) treated with TNF-α and TFGB1 was obtained from the Gene Expression Omnibus under the accession GSE147405 (36). This dataset consists of gene expression values (at the single-cell level) for five different time points (8h, 1 day, 3 days, and 7 days) with treatment and three additional time points (8h, 1 day and 3 day) without treatment, following the last treatment point (7 days). The count values were normalised at each cell level by the total counts over all genes. For each PSMB gene, we first computed the average value across the single cells under each condition and time point. Then, the average expression value of CP (mean of PSMB5, PSMB6, and PSMB7) and IP (mean of PSMB8, PSMB9, and PSMB10) was computed for each condition and time point. The relative change in the expression of IP and CP has been computed with respect to the average expression level at the initial time point (*t*=0).

### 4.5 Genetic alterations in proteasome genes

To check if the CP and IP genes were affected by somatic alterations, we considered non-synonymous mutations and copy number alterations (amplification/deletions) in the six PSMB genes. For the mutation analysis, we removed hypermutator samples (>10 mutations/MB) (63). Further, we computed the fraction of samples having the protein affecting mutation in the IP and CP genes. Similarly, we computed the fraction of samples having copy number alteration (amplified or deleted) in each of these genes in all of 33 tumour types.

### 4.6 Survival analysis using Cox proportional hazards model

To assess the association of proteasome (IP or CP) expression with patient survival, we collected the clinical status of samples from the TCGA Pan-Cancer Clinical Data Resource (64) (TCGA-CDR-SupplementalTableS1.xlsx, https://gdc.cancer.gov/ about-data/publications/pancanatlas). The tumour types (THYM, TGCT, PCPG, DLBC, THCA, and KICH) that were not recommended for overall survival analysis or with low number of cases were removed according to the above TCGA study (64). For the remaining tumour types, we performed univariate and multivariate Cox proportional hazard regression model analyses by considering proteasome (IP or CP) expression and other confounders (such as cytotoxic infiltration level, regulatory T-cell infiltration level, exhaustive T-cell expression level, age, sex, tumour stage, tumour mutational burden) as covariates, independently (univariate) and jointly (multivariate), by using lifeline packages written in Python (<u>https://</u> <u>lifelines.readthedocs.io/</u>). The samples that have missing values for any of the above said parameters were not included. The *p*-values computed using the Wald test were subjected to multiple hypothesis testing correction using the Benjamini–Hochberg approach (see Supplementary Table 6).

### 4.7 Immunoproteasome association with ICB response

To investigate the IP expression association with ICB response, we have collated data from nine publicly available studies across eight tumour types: melanoma (*n*=35 and *n*=26 from two independent studies), lung (*n*=16), thymus (*n*=9), breast (*n*=29), bladder (*n*=348), oral (*n*=11), renal (*n*=11) and eso-gastric (*n*=39) (see Supplementary Table 7 for data source, treatment type and response details). The average proteasome score was computed as the average of log2 transformed values of PSMB5, PSMB6, and PSMB7 for CP and PSMB8, PSMB9, and PSMB10 for IP. To compare the trends across multiple studies, the average scores were transformed into a *z*-score for each tumour type/dataset (Supplementary Table 8). Further, a single sample scoring method (65) was used to calculate the enrichment score of 18 immune signatures and then subjected to differential enrichment analysis between responders and non-responders using the GSVA pipeline.

## Acknowledgements

We thank Dr Dimple Notani and members of RS lab for feedback and suggestions. The results shown here are in part based upon data generated by the TCGA Research Network: https://www.cancer.gov/tcga.

## Funding source

This work was supported by the Department of Atomic Energy, Government of India, under Project Identification No. RTI 4006 and intramural funds from NCBS-TIFR. RS acknowledges support from the DBT/Wellcome Trust India Alliance Fellowship (IA/I/ 20/1/504928) and Bioinformatics Centre Grant funded by Department of Biotechnology, India (BT/PR40187/BTIS/137/9/2021).

## Conflict of interests

The authors declare no conflict of interest.

## Authors’ contributions

RK, BD and RS conceived and designed the study. RK and BD performed the analyses and interpreted the results. SS and MKJ contributed for the EMT analyses and interpreted the results. RK, BD and RS wrote the manuscript with input from other authors. All authors read and approved the final manuscript.

## Availability of data and materials

For TCGA cohort: The pre-processed gene expression (Level 3, RSEM_gene_normalized) data was downloaded from the Broad GDAC firebrowse (http://firebrowse.org/, TCGA data version 2016_01_28). The somatic mutations, copy number alterations, purity and clinical annotations were obtained from the TCGA pancancer atlas study (https://gdc.cancer.gov/about-data/publications/ pancanatlas). For GTEx cohort: the preprocessed gene expression (RSEM gene normalized) was obtained from UCSC Xena browser https://xena.ucsc.edu/. The single-cell gene expression data and dataset related to ICB treatment was obtained from the published studies. The reference for each dataset and processed gene expression values were given in the supplementary tables. The codes used for the analyses and generation of figures is given here https://github.com/onkoslab/ <u>immunoproteasome</u>.

## Supplementary Figures

**Supplementary Figure 1:**
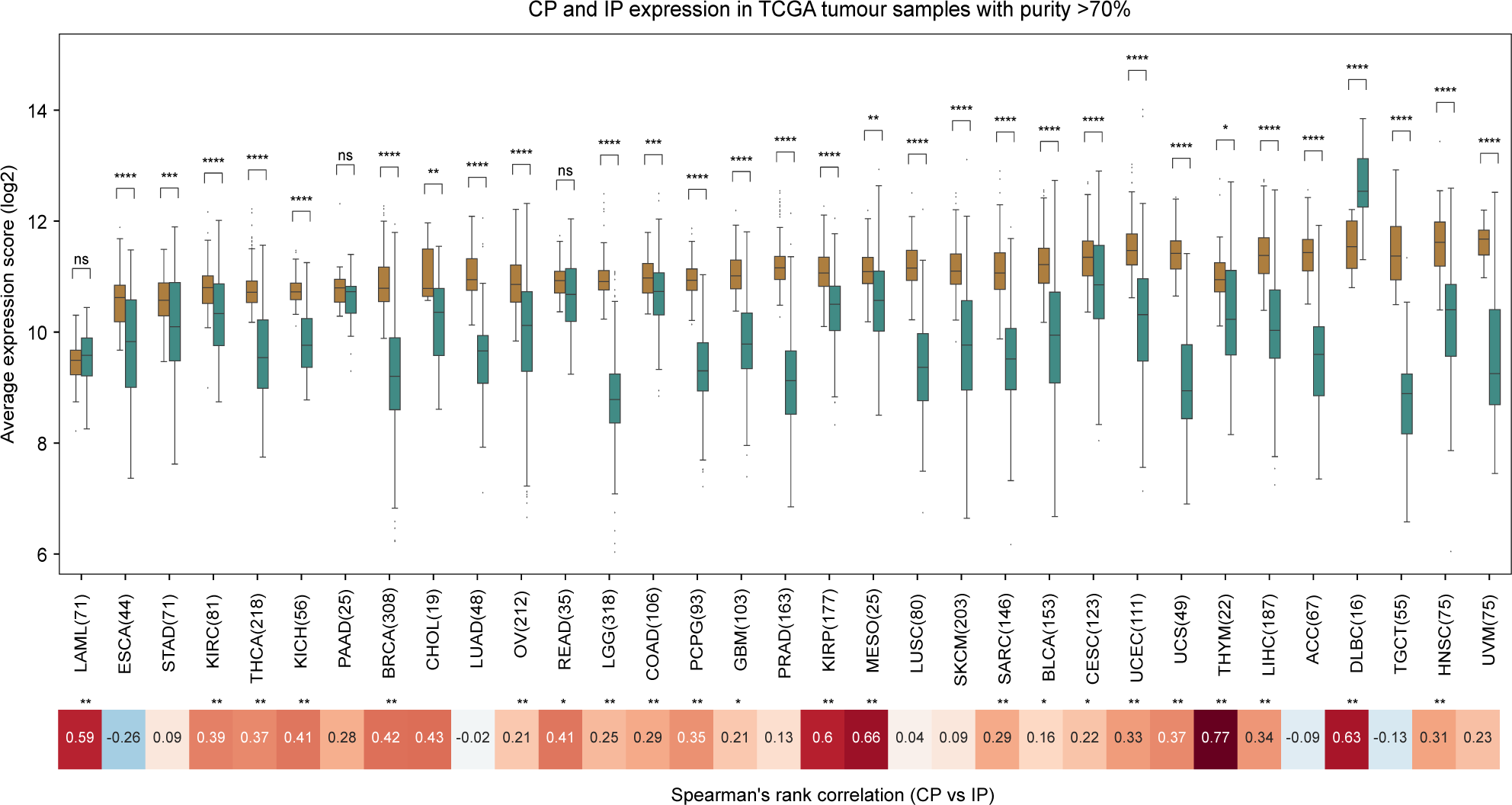
Comparison of CP and IP expression in TCGA tumours with high purity (>70%) The box plot shows the distribution of average expression of constitutive proteasome (CP) and immunoproteasome (IP) genes from 3,535 TCGA samples with high tumour-purity (>70%) across 33 different tumour tissues. The x-axis represents the tumour tissues (with the number of samples that have tumour-purity value >70%) and the y-axis represents the average expression level of CP (PSMB5, PSMB6 and PSMB7) and IP (PSMB8, PSMB9 and PSMB10) genes. The tumour-purity was calculated using the ABSOLUTE algorithm (see Methods). In each boxplot, the horizontal middle line indicates the median, the height of the shaded box indicates the interquartile range (IQR), and the whiskers indicate 1.5 x IQR. The P-value shown at the top, comparing IP and CP expression distributions at each tumour type, was computed using the Mann-Whitney U test (two-sided) and the significance level was represented as: **** P <= 0.0001, *** 0.0001 < P <= 0.001, ** 0.001 < P <= 0.01, * 0.01 < P <= 0.05 or ns - non-significant (P > 0.05). The heatmap at the bottom represents Spearman’s rank correlation between average expression of CP and IP at the sample level for each tumour type. The tumour types that showed significant correlation were highlighted with the asterisks symbol on the top (** P <= 0.01 and * 0.01 < P <= 0.05).

**Supplementary Figure 2:**
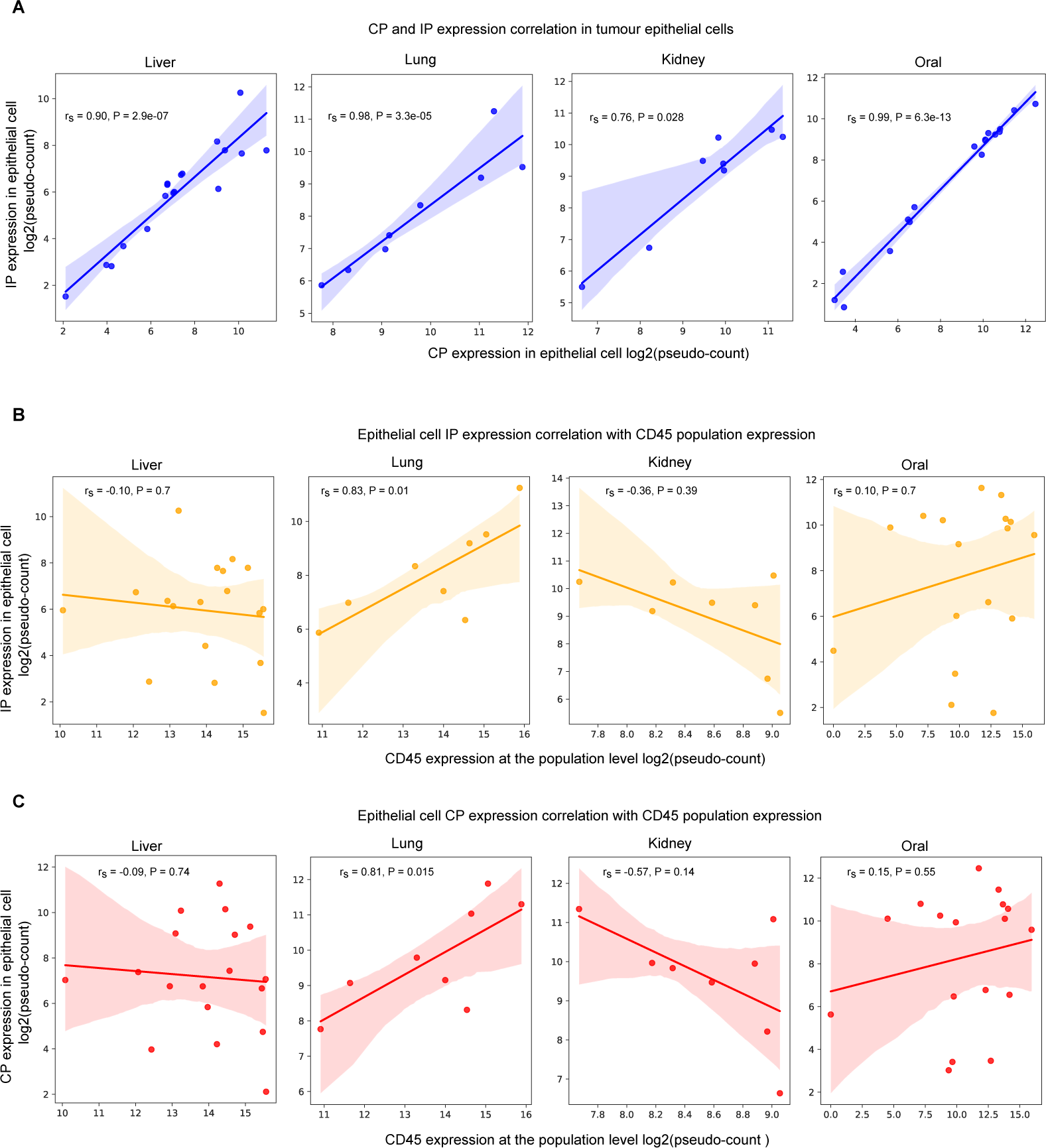
Single-cell analysis of CP and IP expression across tumours. **A.** Spearman’s rank correlation between the average expression of CP and IP genes (from tumour epithelial cell population) in four different cancer tissues (liver, lung, kidney and oral). Each dot on the plot represents a sample and the values on the x- and y-axis represent the CP and IP expression level, respectively, as pseud-count (i.e., average expression of CP or IP genes across all cells within the sample). **B**. Spearman’s rank correlation between CD45 gene expression (from all cells) and IP expression (from epithelial cells) at the sample level in four different cancer tissues (liver, lung, kidney and oral). **C**. Same as B, but for CP.

**Supplementary Figure 3:**
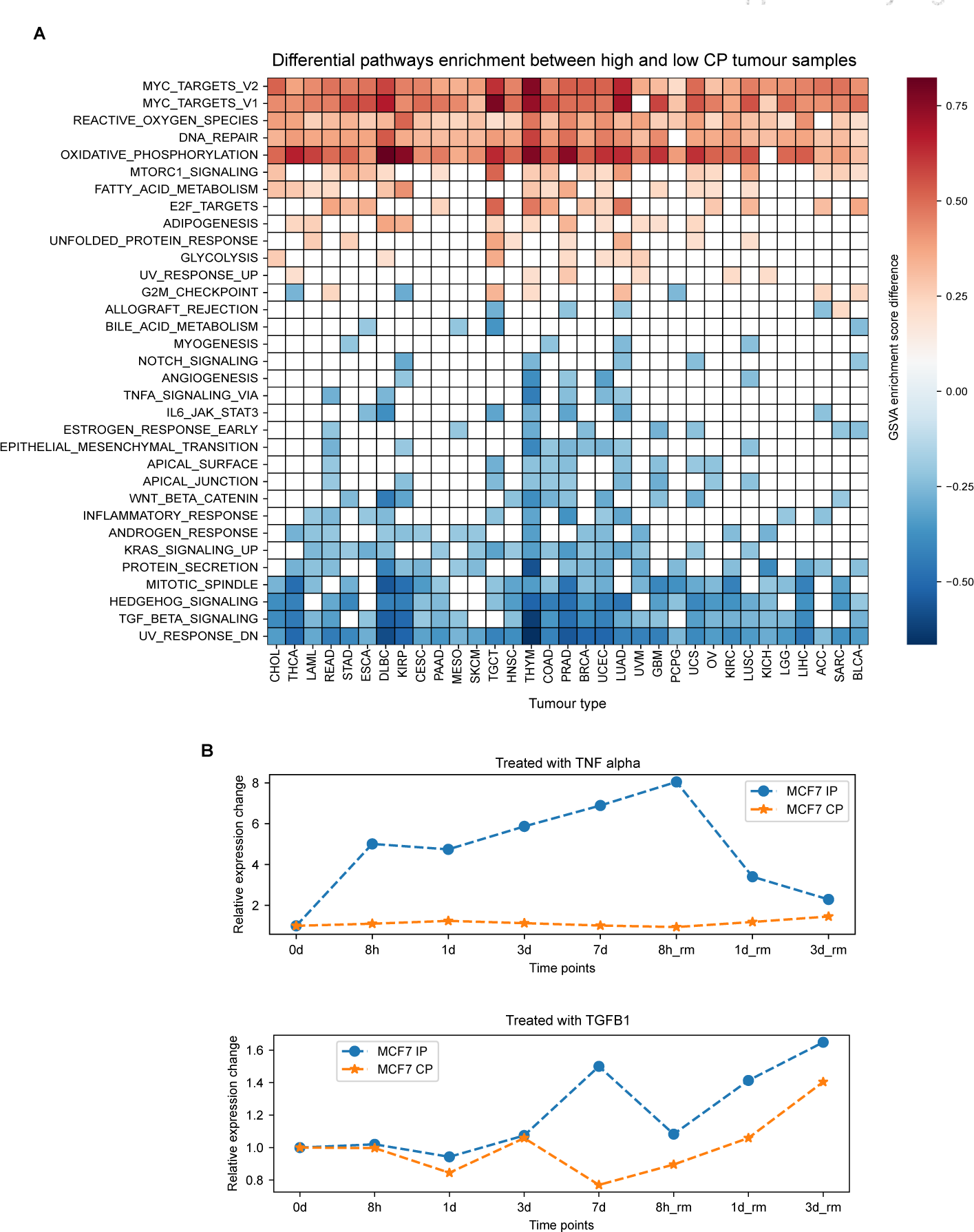
Differential pathway enrichment between high and low CP tumour samples. **A**. Differentially enrichment of hallmark gene-set (from MsigDB) between samples of high-CP (upper quartile) and low-CP (lower quartile) expression groups. The x-axis represents the tumour type and the y-axis represents the different hallmark pathways. The colour of each cell in the heatmap represents the GSVA enrichment score difference between the high-CP and low-CP expression group. Only those pathway that have GSVA enrichment score difference of >|0.2| and FDR adjusted p-value < 0.05 were shown here. The cells with white colour indicate no significant enrichment of pathways in the corresponding tumour, with respect to the above condition. **B.** Relative expression changes in the IP and CP expression level in MCF-7 breast cancer cell line. The x-axis represents the different time points: untreated (0), under treatment (8h, 1 day, 3 days, 7 days) and withdrawal (rm) of treatment after 7 days of treatment (8h, 1 day and 3 days). The y-axis represents the relative expression change with respect to untreated condition (0).

**Supplementary Figure 4:**
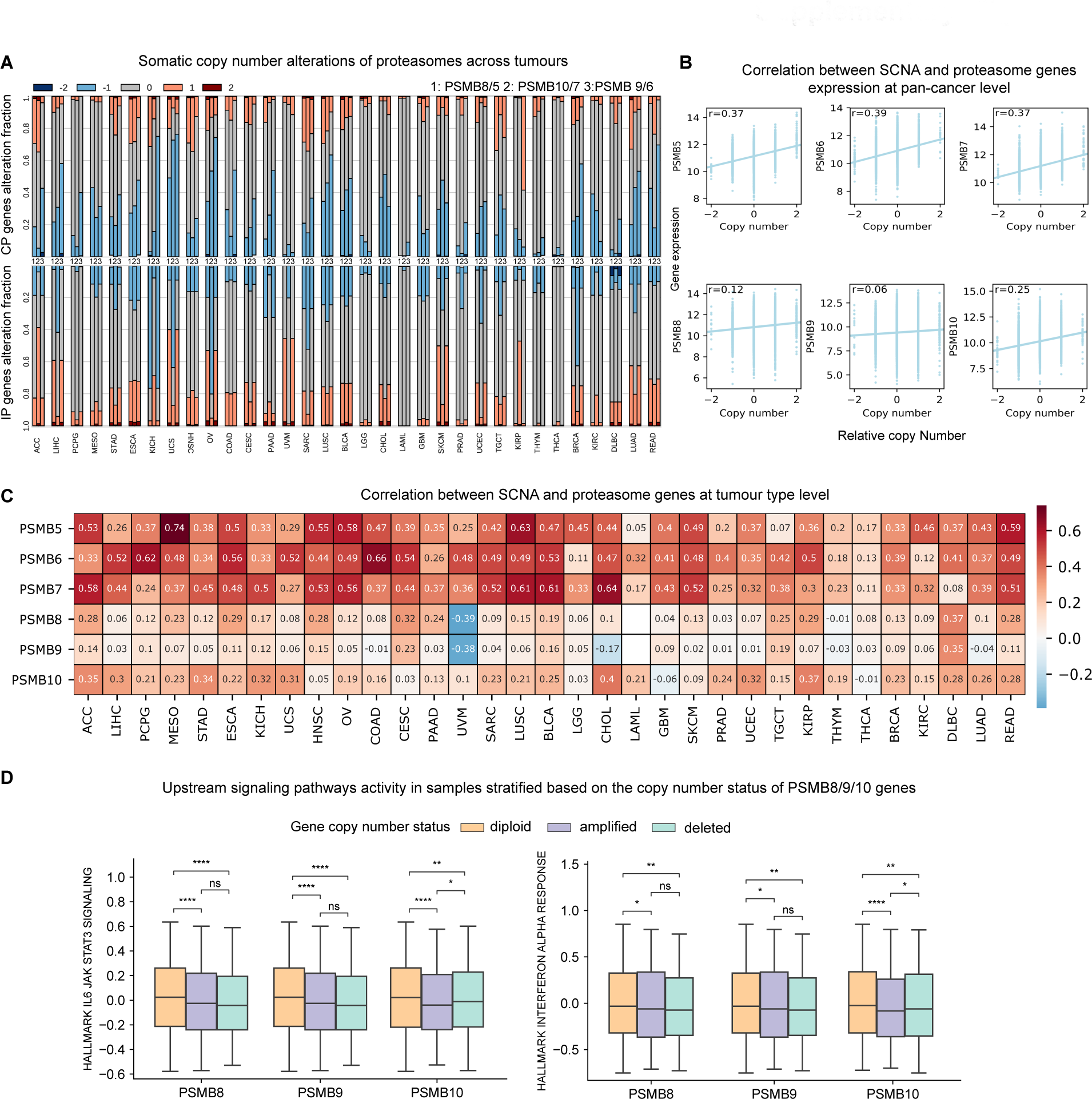
Somatic copy number alterations in proteasome genes and their correlation with gene expression. **A**. Proportion of tumour samples having somatic copy number alteration in CP (PSMB5, PSMB6, PSMB7) and IP(PSMB8, PSMB9, PSMB10) genes. The stacked bar plot represents the fraction of samples with no somatic copy number alterations (zero), deletion (−1), deep deletion (−2), amplification (1) and high level amplification (2) in each tumour type. The top row is for CP genes and the columns 1, 2 and 3 represent PSMB5, PSMB7 and PSMB6, respectively. The bottom row is for IP genes and the columns 1, 2 and 3 represent PSMB8, PSMB10 and PSMB9, respectively. **B-C**.Spearman’s rank correlation between somatic copy number alteration and CP and IP genes individually at the pan-cancer level (**B**), and at the individual tumour type level shown in the bottom heatmap annotated with correlation values. (**C**). **D**. Expression level of IL6_JAK_STAT3 and Interferon alpha response pathways (computed using GSVA) in samples with IP genes copy number altered (amplified CNA>0 or deleted CNA<0) and unaltered (CNA==0, diploid). The p-values were computed using the Mann-Whitney U test (two-sided) and the significance level was represented as: **** P <= 0.0001, *** 0.0001 < P <= 0.001, ** 0.001 < P <= 0.01, * 0.01 < P <= 0.05 or ns - non-significant (P > 0.05).

**Supplementary Figure 5:**
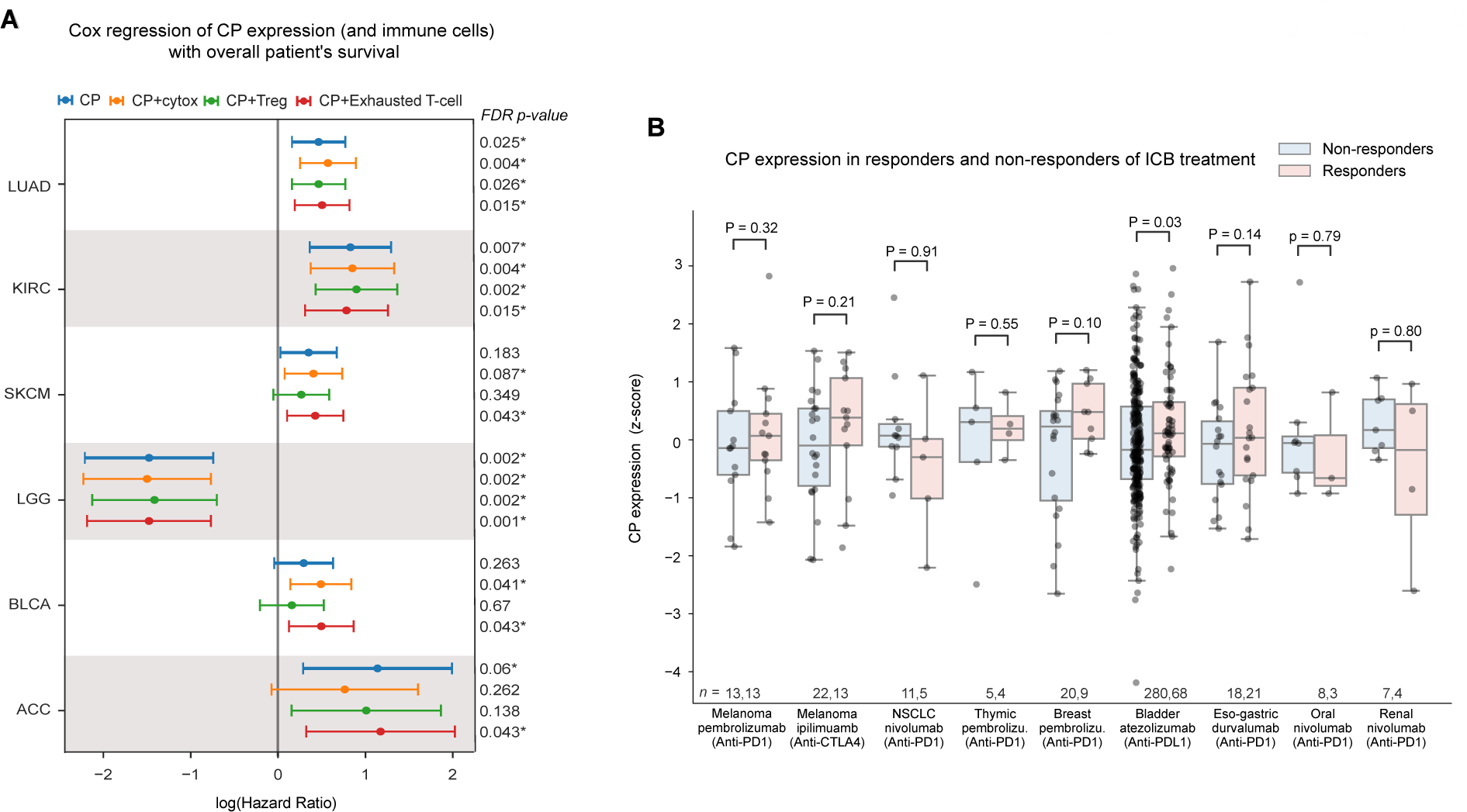
Association of CP expression with overall survival and response to immune checkpoint blockade (ICB) therapies. **A.** Cox proportional hazard ratio (HR) for CP expression alone and in combination with three other confounders (regulatory T cells, Cytotoxic cells and exhausted T cells) in six different tumours from TCGA which showed significant prognostic association. The x-axis represents the log(HR) value (> 0 indicates bad prognosis and < 0 indicates good prognosis), and the horizontal line represents the mean and 95% confidence interval. The colour of the line indicates the variables considered for the analysis. The FDR adjusted p-value shown on the right was calculated using the Wald test and corrected for multiple hypothesis testing using Benjamini-Hochberg method. The asterisks(*) symbol indicates FDR adjusted p-value less than 10% significance level. **B.** The boxplot shows the average CP expression (represented as z-scores) in responders and non-responders of ICB therapies in eight different tumour types. One-sided Mann-Whitney U test was used to compare whether the CP expression in responder was higher than non-responder, and the corresponding P-values were mentioned at the top.

## Supplementary Tables

**Supplementary Table 1:** Gene expression values of IP and CP genes in TCGA and GTEx samples. **A:** Proteasome gene expression level in TCGA samples. **B**: Proteasome gene expression level in GTEx samples. **C**:GTEx sample id matched with the TCGA tumour tissue type. **D**:TCGA sample-id corresponding to low and high expression groups of IP. **E**: TCGA sample-id corresponding to low and high expression groups of CP.

**Supplementary Table 2:** List of different tumour tissues with single-cell transcriptomics data available and their respective references considered in this study.

**Supplementary Table 3:** Sample level GSVA score of 18 different immune cells across 33 tumour tissue samples from TCGA. These scores were further used for the differential enrichment between high and low proteasome (IP or CP) sample groups.

**Supplementary Table 4:** Sample level GSVA score of 50 different pathways across 33 tumour tissue samples from TCGA. These scores were further used for the differential enrichment between high and low proteasome (IP or CP) sample groups.

**Supplementary Table 5:** Proportion of samples having protein affecting mutation in proteasome genes (PSMB5, PSMB6, PSMB7, PSMB8, PSMB9, and PSMB10) at tumour tissue level and at pan-cancer level (last row) in TCGA cohort.

**Supplementary Table 6:** Cox regression analysis of IP (**A**) and CP (**B**) expression with the overall survival in TCGA cohort. The column represents each variable (and its combination) and the values in the row represent log(hazard ratio(HR)) and adjusted p-value. The cells with no value indicate missing values for that variable in the respective tumour type.

**Supplementary Table 7:** Description of dataset related to immune checkpoint blockade (ICB) therapy, with the clinical outcome, and the reference details.

**Supplementary Table 8:** IP and CP expression score in patients samples treated with ICB therapy.

**Supplementary Table 9:** Gene list for 18 different immune cells, 50 different hallmark pathways, epithelial and mesenchymal gene signatures curated from different sources.

## Notes

### Competing Interest Statement

The authors have declared no competing interest.

### Summary of Updates

Figure 3 revised. Methods section updated with more details.

